# Separable multidimensional MRI signatures of cellular and structural pathology in Alzheimer’s disease

**DOI:** 10.64898/2026.04.02.716156

**Authors:** Eppu Manninen, Courtney J. Comrie, Geidy E. Serrano, Thomas G. Beach, Elizabeth B. Hutchinson, Dan Benjamini

## Abstract

Cognitive decline in Alzheimer’s disease (AD) reflects progressive disruption of cellular and microstructural organization, yet the biological specificity of MRI signals remains incompletely understood. Multidimensional diffusion–relaxation MRI (MD-MRI) resolves sub-voxel tissue heterogeneity, offering a potential framework to link imaging signals to underlying pathology. We tested the hypothesis that neuronal, glial and white matter pathologies in AD occupy separable regions of diffusion–relaxation space and generate spatially organized imaging signatures linked to cognitive impairment.

We integrated ex vivo MD-MRI with co-registered histology from 12 human donors spanning a range of Braak stages and pathological severity. Using nested cross-validated elastic net modeling, we predicted voxelwise Aβ, pTau, microglia and myelin burden from the multidimensional diffusion–relaxation density distribution. Regional associations were assessed across hippocampal subfields and white matter, and clinical relevance was evaluated by relating MRI-predicted pathology to Mini-Mental State Examination (MMSE) scores.

Distinct diffusion–relaxation components were preferentially associated with different pathological markers, indicating separable microstructural signatures. Voxelwise MRI-derived predictions were significantly associated with histological measures of myelin (ρ = 0.77), pTau (ρ = 0.62), and microglia (ρ = 0.61), with weaker correspondence for Aβ (ρ = 0.45). Regionally, predicted pathology recapitulated known patterns of selective vulnerability, with elevated pTau and microglial signal in hippocampal subfields and dominant myelin-associated signal in white matter (p < 0.0001). Importantly, higher predicted pTau density in the hippocampus was strongly associated with worse cognitive performance (ρ = −0.88, p = 0.0014), with a moderate association in white matter (ρ = −0.66, p = 0.036), suggesting that tau-related microstructural alterations within both gray and white matter contribute to cognitive impairment.

By directly linking multidimensional MRI signatures to histologically verified cellular pathology, this study demonstrates that AD-related processes manifest as distinct and spatially organized diffusion–relaxation signatures. These findings provide mechanistic insight into the microstructural basis of MRI contrasts and support the potential of MD-MRI to map regionally specific neuropathological processes in AD. As clinically feasible MD-MRI acquisition protocols continue to emerge, translation of these spectral signatures to in vivo imaging may enable more mechanistically informed assessment of aging and dementia.

## Introduction

Alzheimer’s disease (AD) is a progressive neurodegenerative disorder and the leading cause of dementia in older adults, currently affecting more than 7.2 million Americans age 65 and older.^1,2^ Its prevalence is expected to rise substantially as the population ages, intensifying the need for diagnostic strategies that can identify disease processes well before irreversible cognitive decline, particularly because the development and evaluation of disease-modifying therapies depend on detecting pathology at its earliest stages. Although neuroimaging is central to clinical evaluation, existing tools capture only a narrow portion of AD biology. MRI remains primarily sensitive to macroscopic changes such as severe atrophy,^3^ which emerge relatively late, while PET imaging with amyloid-β (Aβ) and phosphorylated Tau (pTau) tracers detects misfolded protein accumulation^4,5^ but offers limited insight into other pathological processes that shape clinical outcomes.^6^ These approaches are therefore blind to the diverse comorbid conditions—ranging from neuroinflammation^7–9^ to oligodendrocytopathy^10–12^—that are present in most individuals with confirmed AD and contribute to its heterogeneous trajectory. Collectively, these limitations underscore the need for imaging methods capable of detecting earlier and broader aspects of AD pathophysiology.

The cellular and molecular landscape of AD extends far beyond Aβ plaques and neurofibrillary tangles (NFTs), even though these lesions continue to anchor diagnostic frameworks such as Braak staging. NFTs follow a predictable pattern that begins in transentorhinal cortex, advances to limbic regions including the hippocampus, and ultimately permeates widespread neocortical areas,^13^ while Aβ accumulates along a distinct neocortical-to-subcortical progression.^14^ Yet these hallmark proteinopathies develop within a dynamic and reactive tissue environment. Increasing evidence highlights the importance of neuroinflammation—particularly microglial and astrocytic activation—which rises in regions of high tau burden and intensifies around plaques and tangles as disease advances.^7–9^ In addition, emerging evidence links early myelin and oligodendrocyte vulnerability to the onset and progression of AD.^11^ As these inflammatory, vascular, and degenerative mechanisms intertwine with classical pathology, the biological underpinnings of cognitive decline become increasingly complex, making it imperative to develop noninvasive imaging biomarkers that can capture this multidimensional landscape noninvasively and enable more accurate assessment of disease stage and progression.

Investigating tissue- and cellular-level changes in the aging brain has been limited by the lack of methods capable of resolving fine structural detail in vivo. Light and electron microscopy provide micrometer- and nanometer-scale resolution but require fixed tissue, restricting their use to postmortem samples in a very narrow field of view. In contrast, MRI enables noninvasive, whole-brain imaging at millimeter-scale resolution—adequate for assessing macroscopic changes such as cortical thickness^15^ or gray matter volume,^16^ but insufficient for direct visualization of cellular-level alterations relevant to aging. Consequently, while recent microstructural MRI studies demonstrate sensitivity to age-related changes, they lack specificity and cannot disentangle the complex meso- and microstructural processes involved.^17–19^ Microscopic tissue features are indirectly reflected in nuclear relaxation rates and water diffusion, which can be probed using quantitative relaxation^20^ and diffusion^21^ MRI. These measures provide coarse estimates of properties such as iron, macromolecular, or myelin content, as well as average cell density and fiber orientation. However, because each voxel encompasses hundreds of thousands of heterogeneous cells and organelles, the relationships between MRI observables and the underlying cellular architecture remain highly ambiguous.

Multidimensional MRI (MD-MRI) has recently emerged as a powerful alternative to conventional diffusion and relaxometry methods that report a single, voxel-averaged metric per contrast.^22–25^ Instead of collapsing the rich microscopic diversity of tissue into mean values, MD-MRI acquires data with simultaneous sensitivity to multiple MR contrasts—typically combinations of diffusion, T_1_, and T_2_—and reconstructs joint distributions of these parameters within each voxel.^26–30^ In this framework, the measured signal is represented as a superposition of distinct water pools, each characterized by its own diffusion and relaxation properties, without imposing a specific biophysical compartment model. This “distribution-based” description offers two key advantages: it mitigates partial-volume effects by revealing multiple components that coexist within a nominal 1–2 mm^3^ voxel, and it leverages the added contrast dimensions to better disentangle different microscopic environments that would be indistinguishable in standard scalar maps. Recent methodological and hardware developments have made such multidimensional correlation experiments practical in biological tissue,^31–34^ enabling preclinical^35^ and clinical^36^ implementations of applications that range from probing microstructural heterogeneity in healthy^37–42^ and diseased^36,43–45^ brain and spinal cord to characterizing pathology in ex vivo human samples.^46–48^ In particular, MD-MRI has been shown to recover signatures consistent with specific cellular and subcellular populations, and to provide effectively intra-voxel microstructural resolution—for example, distinguishing astrogliosis-related changes that remain invisible in conventional diffusion tensor or relaxation imaging.^49^ Taken together, these advances position MD-MRI as a uniquely informative modality for studying complex, heterogeneous tissue, with the potential to improve both mechanistic understanding and in vivo detection of neurodegenerative and inflammatory processes.

Building on these advances, a key unresolved question is how diffusion–relaxation signatures relate to specific neuropathological processes in AD. Here, we sought to test whether diffusion–relaxation signatures reflect specific neuropathological processes and whether they can predict voxelwise Aβ, pTau, microglia, and myelin levels in the human hippocampus. We hypothesized that distinct AD pathologies occupy separable regions of diffusion–relaxation space and give rise to spatially organized imaging signatures. To address this, we adopted a multimodal approach integrating ex vivo MD-MRI with co-registered histology from 12 human donors spanning a range of Braak stages and pathological severity. Using regularized voxelwise modeling, we linked multidimensional diffusion–relaxation features to measured histological burden. By directly relating MRI signals to cellular pathology, this study provides insight into how AD-related pathological processes shape diffusion–relaxation contrast and supports its potential for in vivo application.

## Materials and methods

### Donor specimens

All brain specimens were obtained from adult donors who provided informed consent in compliance with the Common Rule, with all associated health data protected under the Health Insurance Portability and Accountability Act of 1996 (HIPAA).^50^ Twelve temporal lobe samples were provided and processed by the Arizona Study of Aging and Neurodegenerative Disorders/Banner Sun Health Brain and Body Donation Program using their standardized autopsy and postmortem tissue preparation procedures.^50^ Neuropathological evaluation was performed using standardized scoring systems that define the principal pathological features of Alzheimer’s disease, consistent with the National Institute on Aging–Alzheimer’s Association (NIA-AA) criteria for Alzheimer’s disease neuropathologic change.^51^ Each of the twelve samples underwent Braak staging and assessment of Aβ plaques and tau tangles by a certified neuropathologist following established histopathological methods. Tissue sections were taken from the block used for MRI as well as from other brain regions as needed for Braak staging. The density of plaques and tangles in the CA1 subfield of the hippocampus was rated on a four-point semi-quantitative scale (none, sparse, moderate, frequent), which was converted to numerical values ranging from 0 to 3.^52^ Donor demographic, clinical, and pathological information, including Alzheimer’s disease scores, is summarized in Table 1.

**Table 1.** Demographic and neuropathology data for 12 post-mortem samples.

| Subject ID | Age | Sex | PMI (hr) | MMSE | Plaque HP | Tangle HP | Braak stage | CERAD score | NIA-AA ADNC |
| --- | --- | --- | --- | --- | --- | --- | --- | --- | --- |
| 1 | 83 | M | 18.33 | 19 | 1 | 3 | IV | probable | intermediate |
| 2 | 87 | M | 45 | 18 | 1.5 | 0.5 | II | probable | low |
| 3 | 85 | M | 13 | 14 | 3 | 3 | VI | definite | high |
| 4 | 86 | M | 4.75 | 30 | 0 | 1.5 | III | not met | not AD |
| 5 | 75 | M | 5 | 6 | 1.5 | 3 | VI | definite | high |
| 6 | 85 | F | 7.66 | 10 | 3 | 3 | VI | definite | intermediate |
| 7 | 50 | F | 18.5 | 0 | 3 | 3 | VI | definite | high |
| 8 | 89 | M | 10.45 | 26 | 1.5 | 2 | IV | definite | intermediate |
| 9 | 89 | M | 50.5 | 6 | 2 | 2 | V | definite | high |
| 10 | 84 | M | 7.2 | 14 | 2.5 | 3 | V | definite | intermediate |
| 11 | 79 | F | 26 | 15 | 1.5 | 3 | VI | definite | high |
| 12 | 82 | M | 12.83 | 24 | 3 | 1 | III | definite | intermediate |

Post mortem interval (PMI); Mini-mental state examination (MMSE); Hippocampus (HP); Alzheimer's Disease Neuropathologic Change (ADNC).

Following the standardized tissue handling procedures of the Banner Brain and Body Donation Program,^50^ coronal brain sections approximately 1 cm thick were prepared immediately after removal of the brain. The sections were immersed in a 4% formaldehyde solution (commercial formalin) for two days to ensure proper fixation, then transferred to phosphate-buffered saline containing 0.1% sodium azide for storage. Maintaining this uniform preparation protocol and minimizing the postmortem interval helped limit artifacts related to tissue degradation, fixation variability, and rehydration effects.^53^ From each temporal lobe sample, sections were collected for neuropathological staining as outlined previously, while the remaining tissue was reserved for MRI analysis. Before scanning, the samples were carefully trimmed to a maximum diameter of 2.5 cm to fit the MRI coil, ensuring that both the hippocampus and entorhinal cortex were included.

The tissue samples were transferred to the National Institute on Aging (NIA) for imaging experiments conducted as part of this study. At the NIA, each specimen underwent MD-MRI scanning according to the study protocol. Following image acquisition, the samples were shipped back, where they were processed for histological analysis as detailed in the subsequent section.

### MRI acquisition

Upon arrival at the NIA, and before MRI acquisition, each formalin-fixed brain sample was immersed in phosphate-buffered saline (PBS) for approximately 12 days to remove residual fixative and equilibrate tissue hydration. Following this rinsing step, specimens were placed inside 30 mm tubes and submerged in perfluoropolyether (Fomblin LC/8, Solvay Solexis, Italy), an inert, proton-free liquid that produces no background MRI signal. Imaging was performed on a 9.4 T Bruker vertical-bore MRI system equipped with a microimaging probe and a 30 mm quadrature radiofrequency coil.

Multidimensional diffusion–T_2_ measurements were obtained using a three-dimensional diffusion-weighted sequence with a repetition time of 800 ms and isotropic spatial resolution of 200 µm. To sample the multidimensional relaxation–diffusion space (T_2_–D), we applied a hierarchical acquisition scheme as described previously.^47,49^ The T_2_ dimension was encoded by varying the echo time from 11.9 ms to 125 ms, while diffusion weighting was modulated by adjusting the b-value between 2400 s/mm² and 14,400 s/mm² across 13 independent diffusion directions,^54^ using gradient durations of 4 ms and diffusion times of 15 ms. A total of 322 volumes were acquired to construct voxelwise T_2_–D distributions. Comprehensive acquisition parameters are provided in the Supplementary Information.

Additionally, a high-resolution anatomical dataset with 100 µm isotropic voxels was collected using a fast low-angle shot (FLASH) sequence^55^ with a 49.6° flip angle. This dataset served as a structural reference for aligning MRI and histological images.

### Histology and Immunohistochemistry

#### Tissue processing

For immunohistochemistry, AT8 antibody (Fisher Healthcare) and 6E10 (BioLegend) were used at a dilution of 1:1000 to detect pTau and amyloid, respectively. IBA1 (Wako) was used to label microglia at a dilution of 1:5000. Myelin staining was performed using Eriochrome Cyanine (EC).

#### Quantification of histopathology

All histology sections were scanned at 10x magnification using an Olympus VS200 Slide scanner. Quantitative assessment of the microscopy images was achieved through a multi-step image processing workflow. First, digital deconvolution was conducted in QuPath^56^ on each scanned section to isolate the individual staining components—Aβ, pTau, microglia, and myelin—from the background signal, effectively separating them into independent image channels. Once marker-specific images were obtained, optical density values were computed for each stain to quantify labeling intensity. To minimize background noise and exclude artifacts, a thresholding step tailored to each tissue section was then applied. This process ensured that only specific labeling was retained, allowing accurate estimation of the proportion of tissue area positive for each histological marker. An overview of the workflow is shown in Fig. 1A.

**Figure 1.**
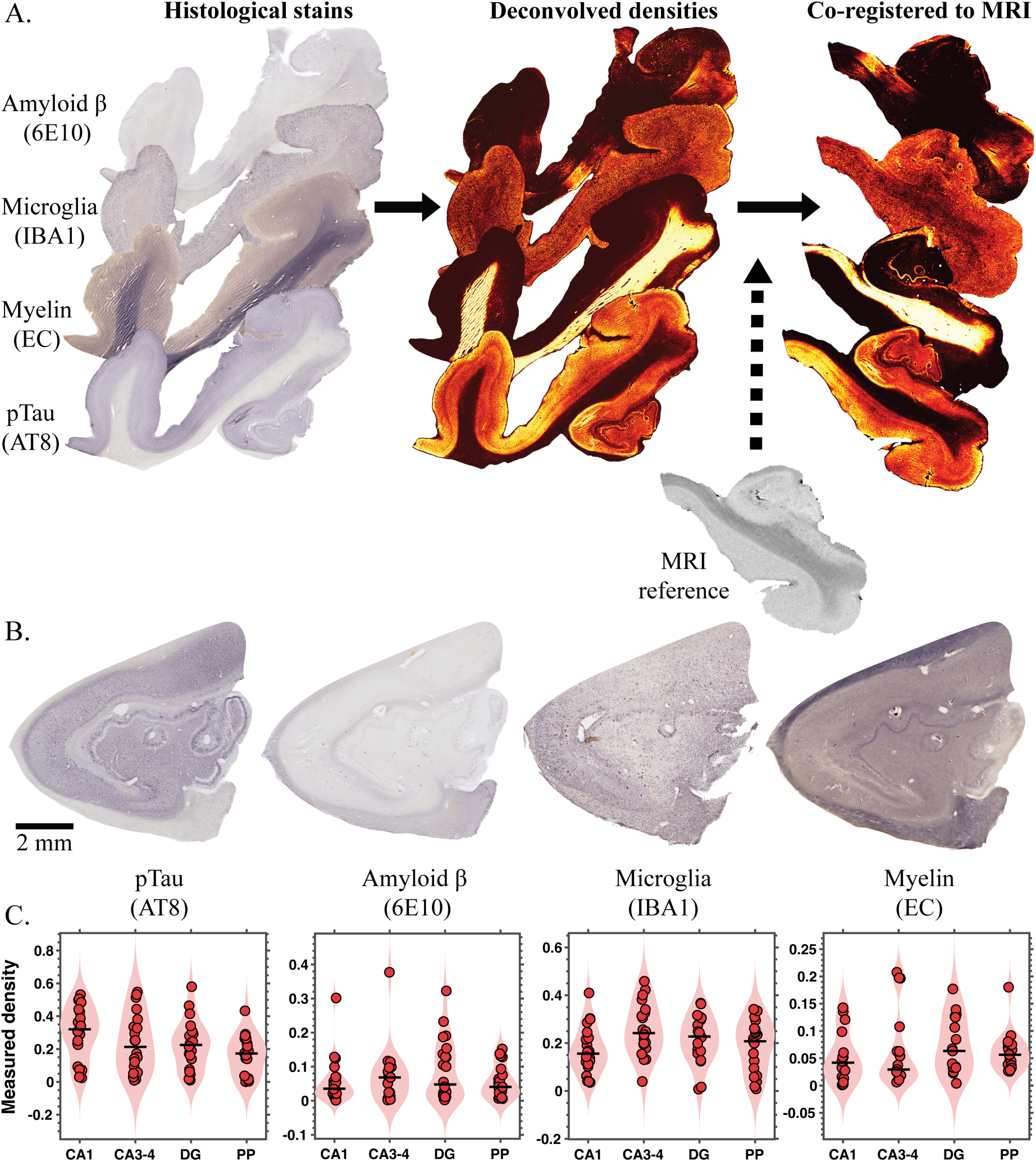
Multimodal registration pipeline and histological findings. **(A)** Four histological stains were first deconvolved to obtain quantitative density maps and subsequently co-registered to high-resolution MRI for voxelwise multimodal alignment. **(B)** Representative pTau, Aβ, microglia, and myelin stains from a hippocampal specimen, illustrating marker-specific spatial patterns and variability in pathological severity. **(C)** Quantified histological burden across cornu ammonis (CA) 1, CA3–4, dentate gyrus (DG), and perforant path (PP) regions for all subjects, demonstrating substantial inter-subject heterogeneity despite generally high neuropathological staging and clinical dementia.

**Figure 1.**
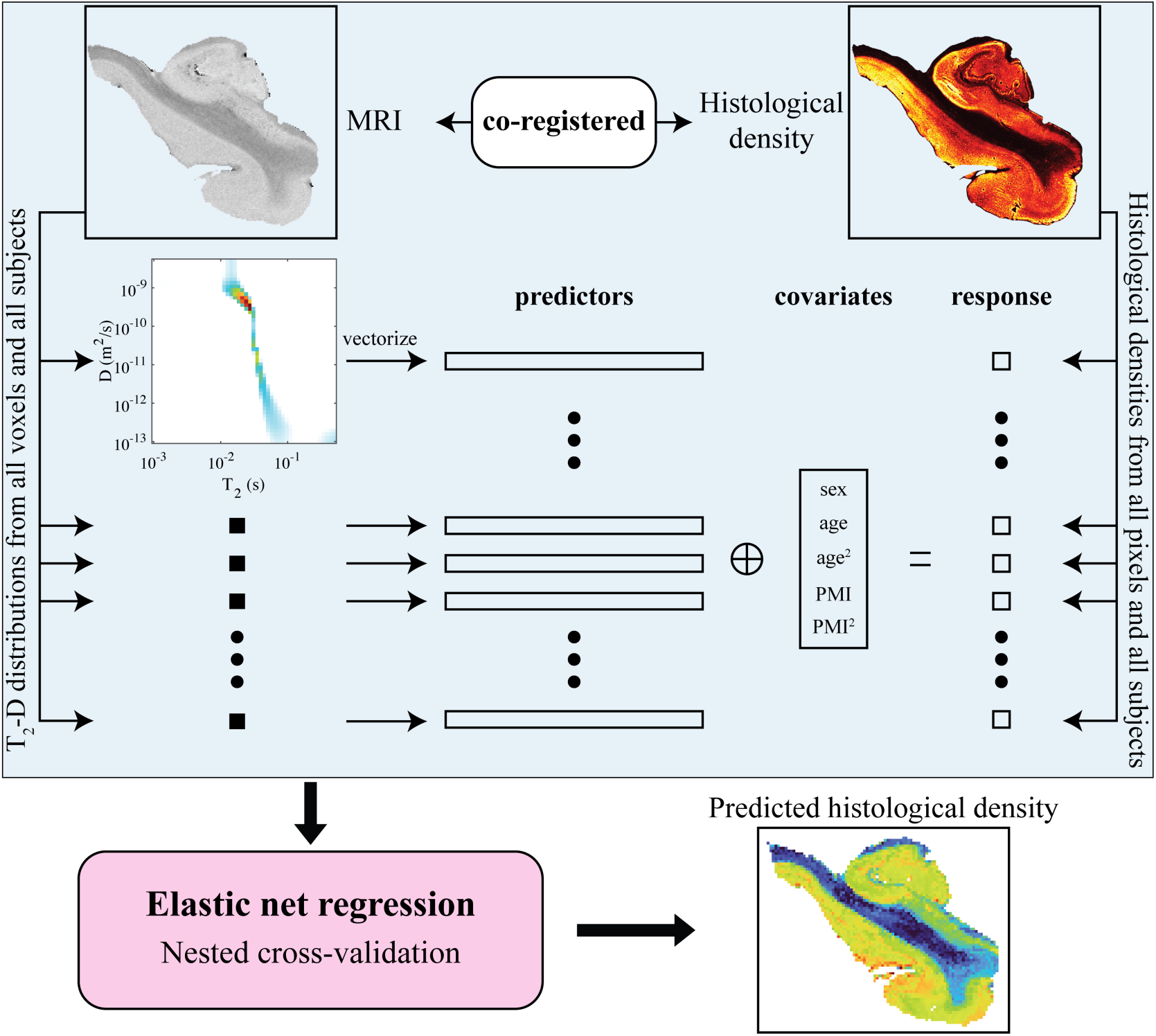
Schematic representation of the proposed framework to predict voxelwise histological outcome from T_2_-D MRI. **(A)** The brain samples were scanned with 3D coverage using MRI. **(B)** Slices from the brain samples were sectioned for histological staining. The acquired histological slices were down-sampled and registered to corresponding MRI slices to allow voxel-wise comparisons between MRI and histology. **(C)** At each voxel, the components of the joint T_2_-D density distribution, along with the biological covariates age, sex and post-mortem interval, were used as predictor variables to model the histological intensity using elastic net-regularized linear regression.

### Registration of histological images to MRI

High-resolution magnetic resonance images served as anatomical references to align the corresponding histological sections. Regions within the histology slides that exhibited significant warping or distortion compared to the original MR anatomy were manually omitted to maintain correct spatial proportions. Following this, alignment between modalities was first performed using two-dimensional affine registration within MATLAB’s Image Processing Toolbox. To achieve finer geometric correspondence and correct residual discrepancies, the affine transformation was refined using a diffeomorphic registration approach in two dimensions. This step employed the publicly available software Greedy (https://github.com/pyushkevich/greedy), which implements the greedy diffeomorphic algorithm,^57^ and was executed following established parameter guidelines from prior literature.^58^ The accuracy of the resulting co-registration was visually confirmed by overlaying the transformed histology on the corresponding MR images. A visualization of the process is shown in Fig. 1A. In total, the dataset comprised 113 histological sections—immunostained for Aβ, pTau, microglia, and myelin—across twelve human brain donors.

### MRI processing

#### Estimation of T2-D distributions

Prior to data analysis, the MD-MRI datasets were denoised using an adaptive nonlocal multispectral filtering algorithm, as previously described.^59^ This preprocessing step enhanced signal quality by reducing noise while preserving spatial and spectral details. Following denoising, voxelwise estimation of the multidimensional relaxation–diffusion distribution was performed using a marginally constrained, nonnegative least-squares optimization method with L2 regularization, in accordance with established procedures.^34,47^ This reconstruction framework, which has been demonstrated in prior studies to provide stable and reproducible results across varying experimental conditions,^60–63^ produced high-fidelity T_2_–D distributions for every voxel in the dataset.

#### Harmonization of distributions between subjects

Tissue processing steps and differing post-mortem intervals between subjects could lead to shifts in the locations of the T_2_-D distribution peaks.^47^ To correct these shifts, we defined a reference distribution for each subject and aligned the voxel-wise distributions of the subjects with a common reference distribution, as detailed in the Supplementary Information and illustrated in Supplementary Fig. 1.

#### Region-of-interest analysis of 3D virtual histology images

To perform region-of-interest (ROI) analysis of the MRI-predicted pathology burden 3D volumes, we created ROIs for white matter and hippocampus (see Supplementary Methods, and Supplementary Fig. 2). To delineate hippocampal subfields, whole-hippocampus ROIs from a previous study^64^ were manually refined in the native space of a high-resolution T_1_-weighted FLASH image using ITK-SNAP software.^65^ The mean diffusivity map was overlaid to facilitate multi-contrast identification of white and grey matter boundaries. Based on established anatomical landmarks,^66^ subfield masks were then generated for cornu ammonis (CA) 1, CA 3–4, dentate gyrus (DG), and the perforant path (PP), and are shown in Supplementary Fig. 3.

### Statistical analysis

#### Modeling histological images using T2-D MRI

We sought to determine whether the voxelwise T_2_-D distribution could predict corresponding histological burden. To address this question, we leveraged a multimodal MRI–histology dataset with voxelwise co-registration, enabling the fitting of linear models in which the components of the T_2_-D distribution at each voxel served as predictor variables and the histological image intensity at the same voxel served as the response variable. We also included age (linear and quadratic terms), sex (binary), and post-mortem interval (linear and quadratic terms) as covariates. A separate single-response model was fitted for each histological modality, with voxels from all subjects pooled during model fitting (Fig. 2). The multimodal MRI–histology dataset with voxelwise co-registration comprised 30, 30, 30, and 23 tissue sections for Aβ, pTau, microglia, and myelin, respectively.

**Figure 2.**
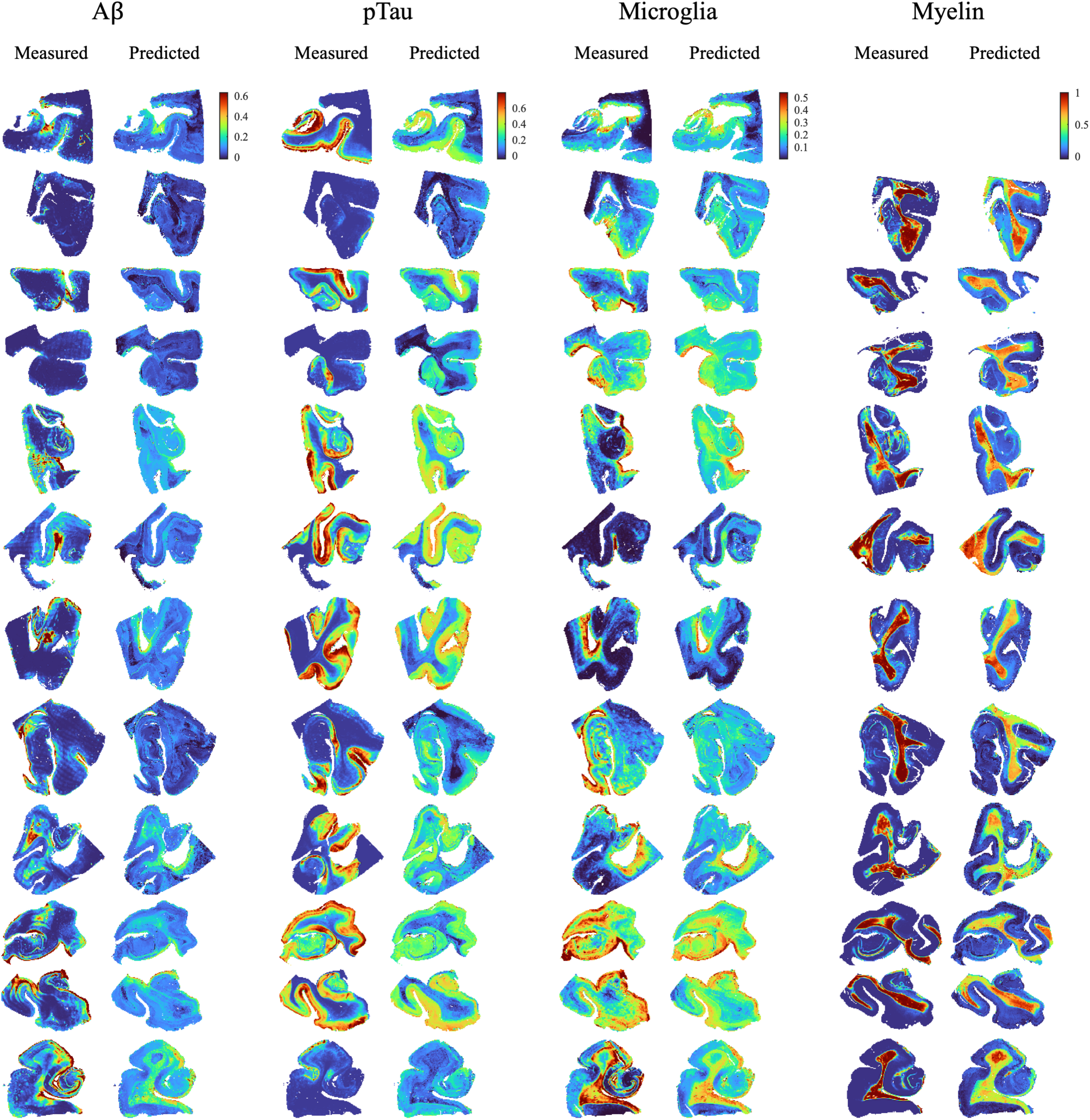
Measured histological and predicted neuropathology burden images. The predicted neuropathology slices were formed by pooling the voxel-wise predictions from the different cross-validation folds. Only one slice shown for each participant. No myelin-stained slices were available for one of the participants. All available slices are shown in Supplementary Figs. 4-7.

The density distribution included many components (50×50=2500) and could have led to overfitted models. To avoid overfitting, we used elastic net regularization.^67,68^ It combines L1 (least absolute shrinkage and selection operator; LASSO) and L2 (ridge) regularization. L1 regularization works as a method for variable selection by promoting sparsity in the solution, tending the coefficients of unnecessary components toward zero. However, if highly correlated predictors exist, L1 regularization tends to select only one and discard the others. This could decrease the performance of the model and lead to incorrect inferences about the importance of the neglected predictors in the model. Adding L2 regularization tends to increase model performance in the presence of highly correlated predictors and improves computational stability by making the optimization problem smoother. We chose equal weighting between L1 and L2 regularization.^69^ R implementation of the software ‘glmnet’ was used for the computations (https://glmnet.stanford.edu/articles/glmnet.html).

As the number of voxels was different for each subject, we added weights to the observations (voxels) so that each subject had the same overall weight in the fitting of the model. To ensure that covariates remained in the model, their penalty factors were set to zero, effectively removing their contribution to the regularization term and preventing their exclusion during model fitting. In contrast, the penalty factors for the T_2_-D distribution components were set to one, allowing them to be subject to regularization. All predictor variables were standardized to unit variance prior to analysis, and an additional non-negativity constraint was imposed on the predictors.

The fit quality of the models was evaluated using nested (externally validated) cross-validation.^70^ This scheme utilizes an outer cross-validation loop to avoid overfitting the hyperparameter (regularization constant) and an inner cross-validation loop to avoid overfitting the predictor variables. To achieve equal representation of voxels from the different slices for each cross-validation fold, we randomly and evenly sampled the voxels from each slice into 10 cross-validation folds. In total, the number of voxels was 189806, 190545, 188937, and 137257 in 30, 30, 30, and 23 co-registered MRI-histology slices of Aβ, pTau, microglia, and myelin, respectively. Because there were 10 cross-validation folds, the training set for each of them contained 90% of the total voxels. We predicted a (cross-validated) response variable (histological intensity) for each voxel and computed the coefficient of determination (R^2^) for each cross-validation fold, along with the root mean squared error (RMSE).

#### Predicting 3D histological burden volumes using MRI

Due to the relatively small number of slices for which histological data is typically available, each model to predict a histological image had to be trained using a maximum of 30 sections divided among the subjects. However, we could use a model trained on those slices to predict the voxel-wise histological responses for whole 3D volumes of voxel-wise T_2_-D distributions. For each set of MRI slices with histological sections available, we used non-nested cross-validation to find the optimal value for the regularization constant and fitted a model with that regularization constant. We then applied that model to the 3D MRI imaging volumes to predict 3D “virtual histology” volumes for each subject. From these 3D volumes, we removed the slices that were contained in the set of training data to avoid training and testing on the same voxels.

#### Regional MRI-derived pathology pattern and dominance analysis

MRI-predicted pathology densities were analyzed across ROIs using non-parametric summary statistics. Regional pathology patterns were characterized by computing the median predicted density across subjects for each ROI and marker, with variability summarized using the interquartile range. Within-marker ROI ranks were additionally computed to facilitate comparison of relative spatial patterning independent of absolute scale.

Then, relative marker dominance within each ROI was assessed using within-subject standardization. Specifically, for each subject and ROI, predicted pathology values were z-scored across markers, such that resulting values represented relative deviation from the within-ROI marker mean. These standardized values were then aggregated across subjects using the median to obtain a robust estimate of consistent relative marker dominance. This approach isolates region-specific differences in relative marker patterns while minimizing the influence of inter-subject variability and global marker scaling.

#### Regional associations between predicted and measured pathology

Associations between MRI-predicted and matched histologically measured neuropathology regional burden were conducted separately for pTau, Aβ, microglia, and myelin, across four hippocampal subregions (CA1, CA3–4, dentate gyrus, and perforant pathway) and WM. One brain sample contained a negligible amount of hippocampal tissue and was excluded from ROI-based analyses. In addition, one subject was substantially younger (50 years) than the remainder of the cohort (75–89 years) and was excluded to minimize age-related confounding. The final analyses therefore included 10 subjects. Given the limited sample size, age and sex were not included as covariates in the ROI-based analyses to avoid overfitting. Within each hippocampal subregion, Pearson correlation coefficients were calculated to assess linear associations between predicted and measured pathology. All correlations were two-sided, and *P*-values of those tests were adjusted using the Benjamini-Hochberg procedure for controlling the false discovery rate (FDR).^71^

#### Associations between predicted neuropathology and cognitive performance

Region of interest-based analyses were also performed to evaluate associations between MRI-predicted pathology measures and mini-mental state examination score (MMSE). ROIs included the whole hippocampus and WM. The same exclusion criteria described above were applied, resulting in a final sample size of 10 subjects. Associations between predicted pathology measures (pTau, microglia, Aβ, and myelin) and MMSE scores were assessed using Spearman rank correlation, given the bounded nature of MMSE scores, the small sample size, and the potential for non-normal distributions. Correlations were computed separately for hippocampal and WM ROIs, and *P*-values were adjusted for multiple comparisons using FDR correction. All tests were two-sided.

## Results

### Neuropathological findings

Table 1 summarizes key demographic characteristics, including latest MMSE and NIA-AA scores. As can be seen, the majority of cases met criteria for dementia at the time of their most recent clinical evaluation. Classic AD neuropathology markers, captured by Braak score and CERAD, ranged in severity. Spearman rank correlation analyses revealed no statistically significant association between MMSE and histologically measured regional plaque burden (ρ = −0.39, *p* = 0.27) or neurofibrillary tangle burden (ρ = −0.60, *p* = 0.13).

Figure 1B illustrates representative pTau, Aβ, microglia, and myelin stains from a hippocampal specimen, highlighting variability in both spatial distribution and severity across markers. For each subject, histological burden was quantified directly from the digitized sections in the study within the CA1, CA3–4, DG, and PP ROIs and is summarized in Fig. 1C. Although many cases had high NIA-AA scores and were diagnosed with dementia, the histologically derived neuropathological measures reveal substantial variability between individuals.

### Predicted neuropathological density maps

The fitted elastic net-regularized linear models were used to generate MRI-derived predicted neuropathology slices by pooling the voxelwise predictions from the cross-validation folds within a nested cross-validation framework. Figure 3 shows a single histologically measured and the corresponding MRI-predicted neuropathological burden slice for each participant and stain. Supplementary Figs. 4–7 show the complete set of slices analyzed across all participants (i.e., total of 113).

**Figure 3.**
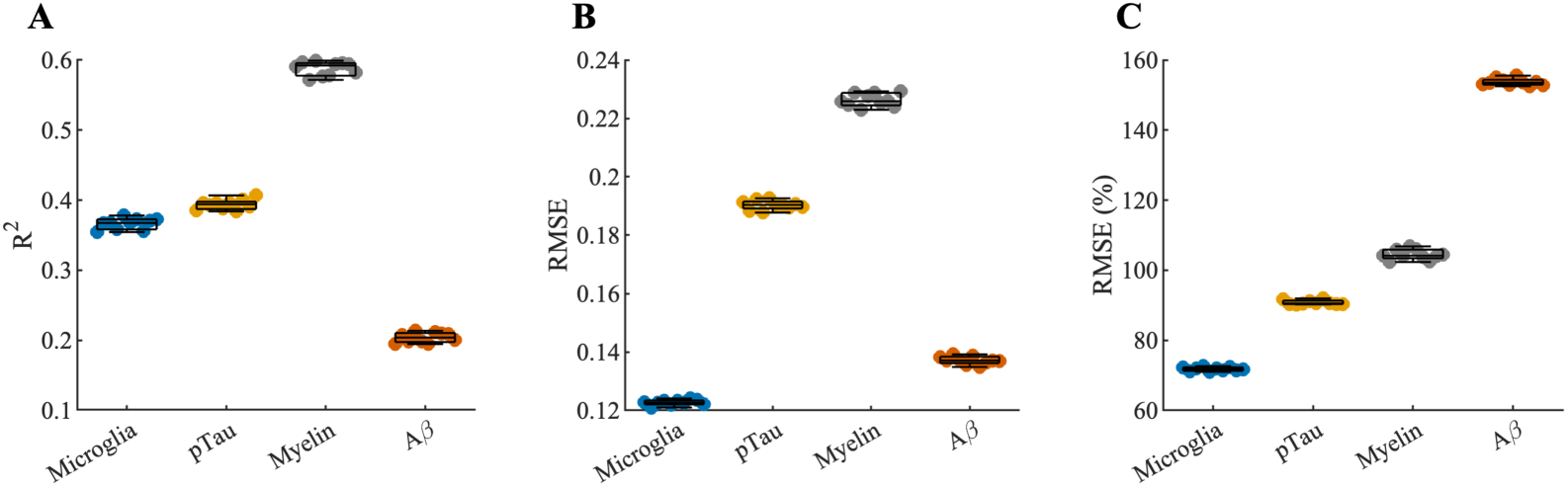
Fit quality metrics of models for predicting histological burden images from T_2_-D distribution images. **(A)** Coefficient of determination (R^2^). **(B)** Root mean squared error (RMSE). **(C)** Relative RMSE. We computed each quality of fit measure using a nested cross-validation framework. Each dot represents the result for one cross-validation fold.

Goodness of fit of the models was assessed by computing the coefficient of determination (*R*^2^), the root mean squared error (RMSE), and the relative root mean squared error (RMSE %; relative to the measured histological values) within a nested cross-validation framework (Fig. 4). After averaging across cross-validation folds, voxelwise associations between predicted and measured neuropathological burden varied by modality. Microglia showed a moderate association (R^2^ = 0.37; Pearson ρ = 0.61), with an RMSE of 0.12 (relative RMSE, 71.8%). Similarly, pTau exhibited a moderate association (R^2^ = 0.39; ρ = 0.62), with an RMSE of 0.19 (relative RMSE, 90.8%). In contrast, myelin demonstrated a stronger association (R^2^ = 0.59; ρ = 0.77), with an RMSE of 0.23 (relative RMSE, 104%). Aβ showed a weaker association (R^2^ = 0.20; ρ = 0.45), with an RMSE of 0.14 (relative RMSE, 154%).

**Figure 4.**
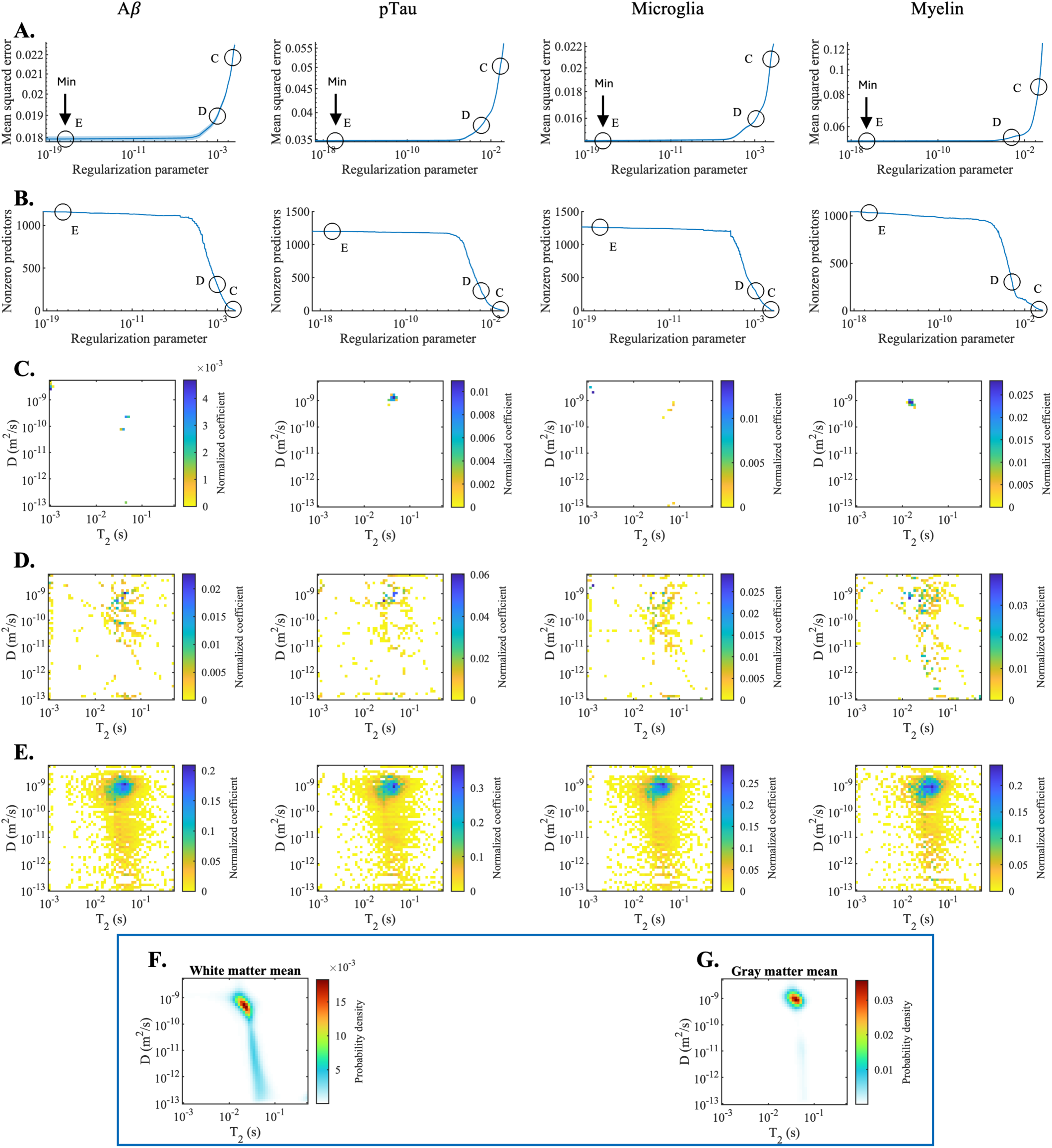
The dependence of neuropathology on diffusion–relaxation features. **(A)** Mean squared error and **(B)** number of nonzero predictor coefficients as a function of the regularization parameter for Aβ, pTau, microglia, and myelin models. Increasing regularization yields progressively sparser models with increasing prediction error. Points C (highly sparse; ∼15 components), D (intermediate; ∼300 components), and E (minimum error) are highlighted. The corresponding normalized fitted coefficients in T_2_–D space are shown in **(C)**, **(D)**, and **(E)**, respectively. The sparsest models **(C)** isolate the dominant diffusion–relaxation feature for each marker, whereas less regularized models incorporate broader regions of T_2_–D space. Color indicates coefficient magnitude. Mean T₂–D distributions across white matter **(F)** and gray matter **(G)** voxels for anatomical reference.

### Neuropathology dependence on diffusion–relaxation features

We next examined which components of the T_2_–D distribution contributed most strongly to the voxelwise prediction models by analyzing the normalized fitted predictor coefficients. Figures 5A and 5B show the mean squared error and the number of nonzero coefficients in the model, respectively, as a function of the regularization parameter. Increasing regularization resulted in progressively sparser models, with a corresponding trade-off between model complexity and prediction error.

**Figure 5.**
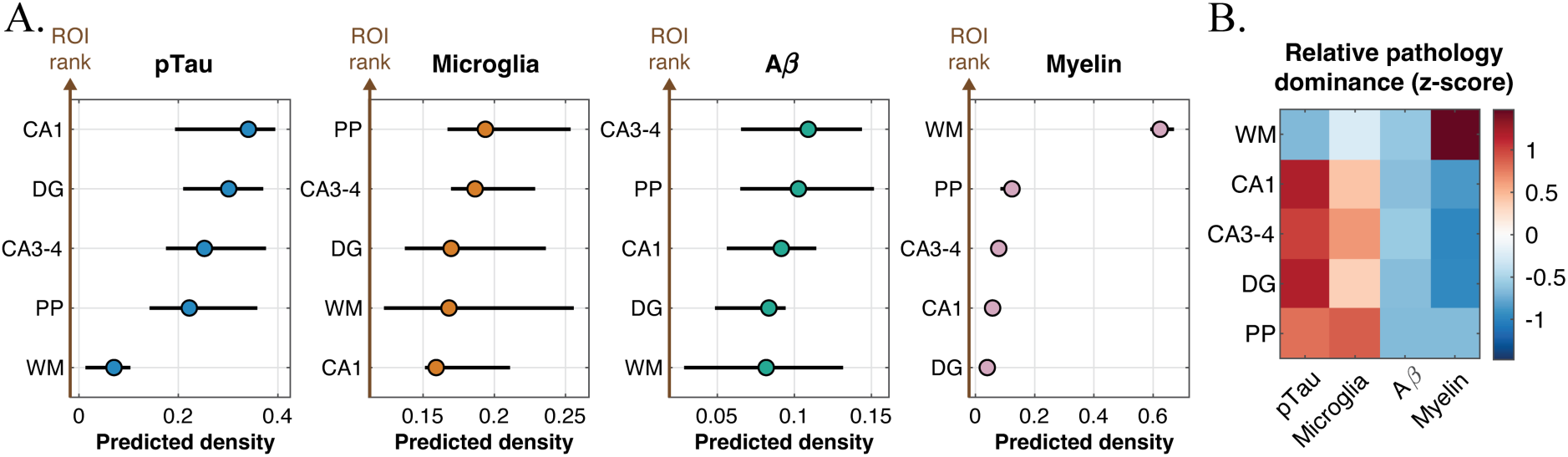
Regional pathology dominance patterns. **(A)** Predicted pathology densities across white matter (WM) and hippocampal subfields that included cornu ammonis (CA) 1, CA3–4, dentate gyrus (DG), and perforant path (PP), for pTau, microglia, Aβ, and myelin. Points indicate mean predicted density; error bars denote variability across subjects. pTau and microglial signals were elevated in hippocampal subfields relative to WM, whereas myelin-associated signal was highest in WM. Aβ predictions showed weaker regional differentiation. ROIs are displayed in descending order of predicted density for each marker. **(B)** Relative pathology dominance across ROIs, calculated using within-subject, within-ROI z-scoring across markers. Warmer colors indicate relative dominance and cooler colors relative suppression. Hippocampal ROIs preferentially exhibited pTau and microglial dominance, whereas WM showed strong myelin dominance. These patterns were conserved across subjects and were independent of absolute pathology magnitude.

Three representative points along this regularization path were selected for further inspection: point C, corresponding to a highly sparse model with approximately 15 nonzero components; point D, corresponding to an intermediate model with approximately 300 nonzero components; and point E, corresponding to the model with the lowest mean squared error (i.e., the optimal model that was used for subsequent computations). The spatial distribution and magnitude of the fitted coefficients for these three operating points are shown in Fig. 5C–E.

Across models, T_2_–D components with larger coefficient magnitudes and those that remained nonzero under stronger regularization contributed most consistently to histological prediction, indicating that these components capture robust features of tissue microstructure. The sparsest models (Fig. 5C) were particularly informative, as they isolate the single most salient diffusion–relaxation feature associated with each neuropathological marker.

Using these sparse models, we computed the weighted average [T_2_, D] coordinates of the retained components, yielding values of [24.6 ms, 1.40 µm²/ms] for Aβ, [43.1 ms, 1.28 µm²/ms] for pTau, [21.7 ms, 1.83 µm²/ms] for microglia, and [15.6 ms, 0.80 µm²/ms] for myelin. Several distinctions emerged across markers: myelin was associated with the fastest T_2_ relaxation and lowest diffusivity, microglial signal with the highest diffusivity, and pTau with the slowest relaxation. For anatomical reference, Figs. 5F and 5G show the average T_2_–D distributions across all white matter and gray matter voxels, respectively, providing context for interpreting the T_2_–D components identified by the model.

### Distinct regional pathology dominance patterns

A key advantage of MR-based neuropathology prediction is its reliance on whole-volume tissue information, in contrast to single histological slices with limited fields of view. MRI-predicted pathology densities across the entire 3D volume exhibited distinct, marker-specific spatial patterns across ROIs (Fig. 6A). For pTau and microglial markers, hippocampal subfields (CA1, CA3–4, dentate gyrus, and perforant path) showed consistently elevated predicted densities compared with the other ROIs, while WM demonstrated lower levels. In contrast, myelin-associated pathology showed its highest predicted densities in WM, consistent with its known anatomical distribution, with minimal involvement of hippocampal regions. Aβ predictions displayed a more distributed pattern with weaker regional differentiation.

**Figure 6.**
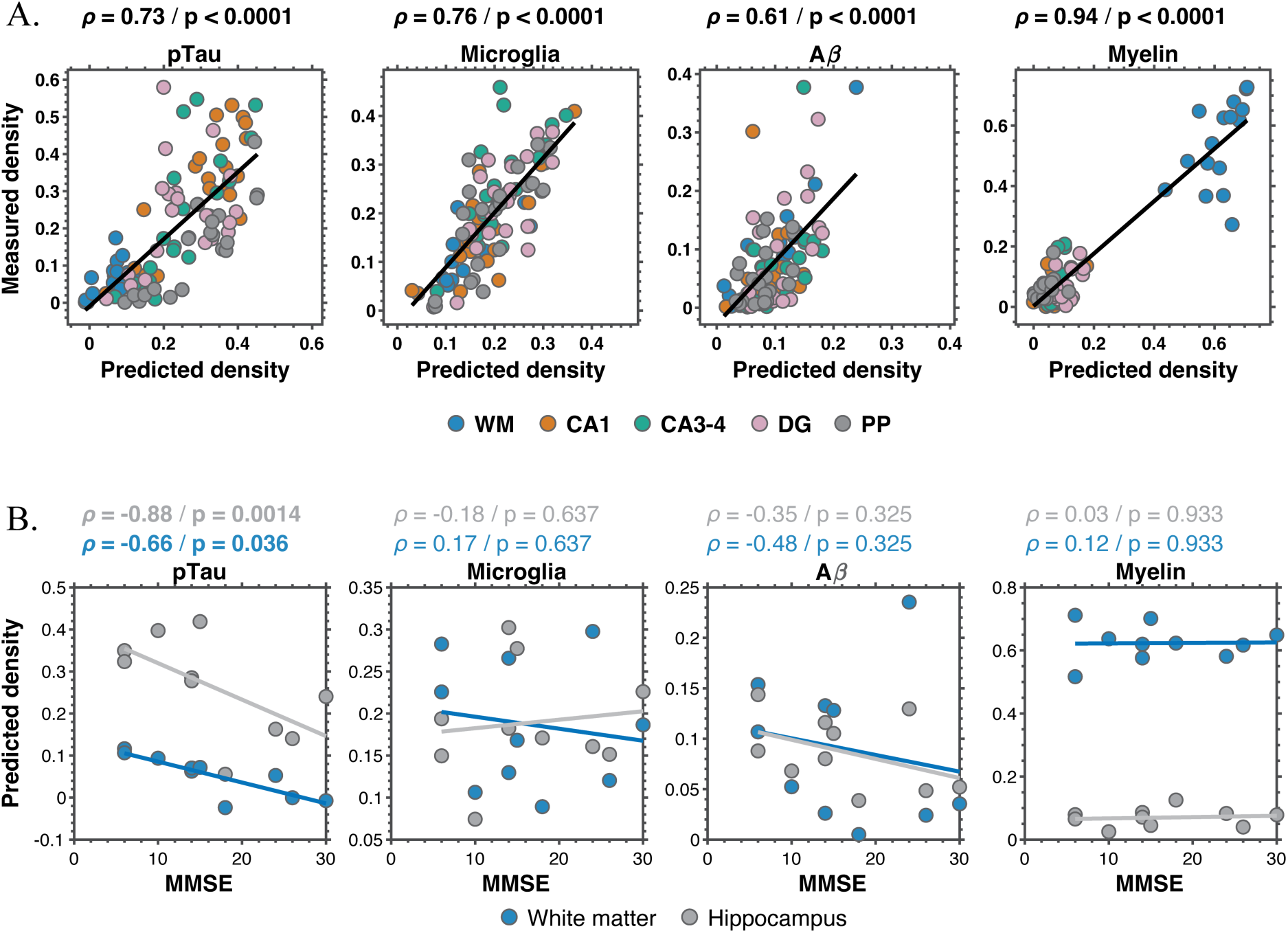
Regional MRI-predicted pathology and cognitive associations. **(A)** Correlation between MRI-predicted and measured histological densities across the cornu ammonis (CA) 1, CA3–4, dentate gyrus (DG), and perforant path (PP) regions of interest for pTau, microglia, Aβ, and myelin. Each point represents a regional measurement; lines indicate linear fits. Robust associations were observed for all markers (p < 0.0001). **(B)** Associations between MRI-predicted regional density and cognitive performance (Mini-mental state examination, MMSE) in whole hippocampus (grey) and white matter (blue). Predicted pTau density showed significant inverse associations with MMSE in both regions, whereas microglial, Aβ, and myelin measures were not significantly related to cognitive performance.

To assess whether these spatial patterns reflect relative regional preferences independent of absolute magnitude, we performed an ROI–neuropathology dominance analysis based on within-subject, within-ROI z-scoring across markers (Fig, 6B). This analysis revealed a clear anatomical stratification of relative pathology dominance. Hippocampal ROIs consistently showed positive dominance of pTau and microglial predicted densities relative to other markers, whereas Aβ was comparatively suppressed. White matter exhibited strong relative dominance of myelin, consistent with its expected tissue specificity and serving as an internal consistency check of the dominance framework. Importantly, these dominance relationships were conserved across subjects and were not driven by overall pathology burden, highlighting region-specific marker preference patterns beyond trivial anatomical effects.

### Regional MRI-predicted pathology aligns with measured histology

To assess regional agreement between MRI-predicted and measured histological burden, all ROIs (WM, CA1, CA3–4, DG, and PP) were pooled and correlated with corresponding histological densities (Fig. 7A). Robust and highly significant associations were observed for all four markers, including pTau (ρ = 0.73, *p* < 0.0001), microglia (ρ = 0.76, *p* < 0.0001), Aβ (ρ = 0.61, *p* < 0.0001), and myelin (ρ = 0.94, *p* < 0.0001).

### MRI-predicted pTau burden is associated with cognitive status

Associations between MRI-predicted volumetric neuropathological density and MMSE were evaluated using Spearman rank correlation in the whole hippocampus and WM ROIs (Fig. 7B). Predicted pTau density showed a strong inverse association with MMSE in the hippocampus (ρ = −0.88, *p* = 0.0014) and a moderate inverse association in white matter (ρ = −0.66, *p* = 0.036), indicating higher tau pathology with greater cognitive impairment.

In contrast, predicted microglial density did not show a clear relationship with MMSE in either ROI. Predicted Aβ density showed a modest inverse trend with MMSE in WM (ρ = −0.48), but this association did not reach statistical significance, and no corresponding trend was observed in the hippocampus. Predicted myelin and microglia densities were not associated with MMSE in either region.

## Discussion

This study demonstrates that multidimensional diffusion–relaxation MRI can capture distinct signatures of AD-related pathology in the human hippocampus when anchored to histopathology. A key strength of this approach lies in its ability to resolve distinct components of the T_2_–D distribution, which serve as spectral representations of tissue microstructure and composition. Leveraging these components, we developed linear models to predict voxelwise Aβ, pTau, microglial, and myelin densities in ex vivo hippocampal tissue from AD donors. Voxelwise MRI-derived predictions were significantly associated with co-registered histological measures of myelin, pTau, and microglial burden, with comparatively weaker correspondence for Aβ. At the regional level, MRI-derived pathology estimates showed robust concordance with measured histological burden across all markers. Extending these findings to clinical relevance, higher MR-predicted pTau burden in both the hippocampus and WM was associated with worse cognitive performance, as reflected by lower MMSE scores.

While Aβ and pTau remain hallmark markers of AD pathology, and cost-effective approaches for measuring them—most notably via plasma—are rapidly advancing,^72,73^ a more holistic characterization of the disease is increasingly needed. Such a broader perspective must account for neuroinflammatory mechanisms, in which microglia play a central role and may exert divergent effects depending on disease stage and individual susceptibility,^74^ as well as the frequent presence of comorbid pathologies that are not captured by biomarker-specific assays. Converging work also highlights myelination as a fundamental component of AD pathophysiology, with emerging evidence suggesting that loss of myelin integrity may represent an upstream risk factor for neuronal Aβ deposition.^12,75^ While T_2_-based MRI approaches for myelin water imaging have gained increasing traction,^76^ and preliminary associations between pTau and diffusion MRI have been reported,^77^ direct estimation of pTau, microglial, and Aβ pathology using MRI—validated through voxelwise co-registration with histology—remains largely unexplored. In this context, our work introduces a unified framework for predicting all four pathological markers and mapping their spatial distributions in the human hippocampus ex vivo.

MRI signals are inherently indirect reflections of tissue microstructure, limiting biological specificity when interpreted in isolation. By integrating ex vivo multidimensional diffusion–relaxation MRI with spatially co-registered high-resolution histology, we established a direct voxelwise framework for linking MRI-derived spectral features to underlying neuropathology. This cross-modal alignment enables quantitative comparison across spatial scales, with microscopy serving as a biologically grounded reference for local tissue composition. The modeling approach is theoretically well matched to the T_2_–D representation, in which each voxel is expressed as a weighted combination of diffusion–relaxation components, allowing interpretable mapping between distinct microstructural signatures and histological burden. Importantly, the analysis was performed voxelwise rather than within predefined ROIs, thereby avoiding spatial averaging and selection bias and permitting radiological–pathological correlations at their native resolution. Although voxelwise effect sizes were more modest than ROI-level associations (e.g., pTau resulted in ρ = 0.62 for voxelwise, ρ = 0.73 for ROI), this is expected given the absence of spatial aggregation and the intrinsic variability of co-registration and histological signal. The combined use of elastic net regularization, subject-balanced weighting, and nested cross-validation further supports the methodological rigor and robustness of this voxelwise MRI–histology mapping framework.

The voxelwise prediction models showed strong ordinal correspondence between MRI-derived predictions and histology for microglia, pTau, and particularly myelin, as reflected by relatively high coefficients of determination, despite limited quantitative agreement as indicated by higher relative RMSE values (Fig. 4). This pattern reflects the fact that histological staining intensity is inherently semi-quantitative and subject to nonlinearities and saturation effects, making a one-to-one quantitative match between MRI-derived predictions and histology neither expected nor required. Rather than reproducing absolute histological values, MD-MRI captures ordinal relationships with underlying pathological burden. In contrast, voxelwise prediction of Aβ was comparatively weaker (Figs. 3–4). One plausible explanation is the highly sparse nature of Aβ neuropathology in the current sample, characterized by a small number of voxels with high plaque burden and many voxels with negligible signal. Under such conditions, even minor mis-registration between MRI and histology can disproportionately affect voxelwise model fitting. In addition, the imbalance introduced by sparse Aβ signal during model training may further limit predictive performance.

The diffusion–relaxation coordinates identified by the sparsest models, which are shown in Fig. 5C, provide mechanistic insight into the microstructural signatures underlying each predicted histological marker. Because these models isolate the single most influential T_2_–D component for each pathology, the corresponding weighted [T_2_, D] coordinates can be interpreted as the dominant diffusion–relaxation signature associated with that tissue feature. Specifically, rapid relaxation and low diffusivity for myelin is consistent with earlier observations^78,79^ and with the principles underlying myelin water fraction imaging. Moreover, the relatively high diffusivity associated with the microglial model (i.e., 1.83 µm²/ms) may reflect the contribution of permeable or exchange-prone membrane structures.^80^ Together, these findings indicate that distinct neuropathological processes occupy separable regions of diffusion–relaxation space.

Consistent with the component-level signatures, the spatial distribution of the predicted maps mirrors known patterns of anatomical vulnerability within the hippocampal formation (Fig. 6A). Predicted pTau and microglial pathology were higher in hippocampal subfields—regions particularly susceptible to neurofibrillary degeneration and inflammatory activation in AD—whereas myelin-associated signal was greatest in WM, in line with its structural composition. The dominance analysis (Fig. 6B) further supports that these differences reflect marker-specific regional involvement rather than uniform scaling of pathology across the tissue. Such regionally differentiated patterns are particularly relevant given the central role of hippocampal integrity in cognitive decline, raising the question of whether these spatially resolved pathology signatures relate to clinical status.

The robust inverse relationship between hippocampal predicted pTau density and MMSE (Fig. 7B) aligns with converging evidence that tau burden, rather than amyloid load, more closely tracks clinical severity in AD.^81,82^ Given that the hippocampus is among the earliest and most severely affected regions in the Braak trajectory, this regional association is consistent with the established link between neurofibrillary pathology and cognitive decline. The association observed in WM is also noteworthy. Although tau pathology is often framed in GM terms, accumulating evidence links tau burden to WM microstructural alterations and structural connectivity changes associated with cognitive impairment.^83–87^ While these findings should be interpreted cautiously given the modest sample size, they are compatible with models in which tau-related network disruption contributes to clinical decline. By contrast, the absence of clear associations between MMSE and predicted microglial or myelin measures suggests that these processes, although biologically important, may not directly map onto global cognitive performance within the hippocampus.

Although diffusion–relaxation MRI showed promise for estimating neuropathological burden in AD post-mortem tissue, several design-related limitations should be considered. The most notable is the relatively small sample size, which reflects the technical challenge of assembling rigorously co-registered multimodal MRI–histology data from the same specimen. Such integration is demanding due to differences in resolution and contrast, tissue deformation during processing, non-coplanar imaging planes, and incomplete or damaged histological sections. While most post-mortem MRI studies include only 5-10 specimens,^46,88–90^ the limited number of cases in the present study nonetheless constrains statistical power, especially in light of the substantial inter-subject neuropathological variability. Several specimens also exhibited neuropathological comorbidities common in aging brains, which may influence diffusion–relaxation MRI signatures and complicate attribution of signal changes solely to AD pathology. However, this highlights a potential advantage of MRI, whose sensitivity to diverse tissue alterations may capture pathology beyond canonical AD biomarkers. At the same time, post-mortem MRI provides an essential intermediary step between histology and in vivo imaging, clarifying the biological basis of multidimensional diffusion–relaxation contrasts. Importantly, recent advances have enabled clinically feasible implementations of MD-MRI protocols,^36,40,41,91^ suggesting that translation of the present findings to living subjects is achievable from an imaging perspective. However, while the current study establishes a mapping between histology and ex vivo MRI, extending this approach in vivo requires an additional step: linking ex vivo and in vivo MRI measurements within the same subject. Establishing this bridge between in vivo imaging and histology remains the principal challenge for clinical implementation. Carefully designed animal models and hybrid ex vivo–in vivo MRI studies may provide a controlled framework for facilitating this transition while enabling accurate voxel-level co-registration.

In summary, by linking multidimensional diffusion–relaxation MRI with co-registered histology in human hippocampal tissue, this study demonstrates that distinct AD pathologies occupy separable regions of diffusion–relaxation space and give rise to spatially organized imaging signatures. The ability to predict voxelwise pTau, microglial, and myelin burden—and to detect associations between tau-related signal and cognitive impairment—indicates that MD-MRI captures biologically meaningful features of selective vulnerability in AD. These findings enhance the interpretability of MRI contrasts and move the field closer to noninvasive mapping of the spatial organization of neuropathology. As multidimensional acquisition protocols become increasingly feasible in clinical settings, translating these spectral signatures into in vivo biomarkers represents a promising step toward mechanistically informed imaging of aging and dementia.

## Data availability

The datasets generated and analyzed during the current study are available from the corresponding author upon request. MATLAB source code for preprocessing and MADCO data inversion is freely available at https://github.com/dan-benjamini/madco/.

## Acknowledgements

This research was supported by the Intramural Research Program of the National Institutes of Health (NIH). The contributions of the NIH author(s) are considered Works of the United States Government. The findings and conclusions presented in this paper are those of the author(s) and do not necessarily reflect the views of the NIH or the U.S. Department of Health and Human Services. We are grateful to the Banner Sun Health Research Institute Brain and Body Donation Program of Sun City, Arizona for the provision of human biological materials. The Comparative Pathology Core Laboratory (CPCL) at the University of Arizona performed all slide scans. We thank Mr. Hiram Cervantes for assistance in developing the subfield ROI segmentation.

## Supplementary material

Supplementary material is available at *Brain* online.

