## Supplementary for "Separable multidimensional MRI signatures of cellular and structural pathology in Alzheimer’s disease"

### Supplementary Methods

#### MRI acquisition

The acquisition of multidimensional data was done using echo planar imaging (EPI) readout according to the MADCO framework encoding scheme,[1,2] and by varying the following two experimental parameters: the echo time, TE, and the diffusion weighting, *b*, providing $T_2$-, and diffusion-weighting, respectively.

The minimal TE value depends on the sample physical dimensions because of the varying imaging matrix size that is intended to keep the spatial resolution constant at 200 µm isotropic voxels. We kept the minimal TE constant across the samples at 11.9 ms using 16 EPI segments.

The two 1D distributions of $T_2$ and MD were estimated, respectively, with the following data acquisition protocols: For $T_2$ encoding, a 1D $T_2$-weighted data set ($b$=0) with 20 logarithmically sampled TE values ranging from 11.9 to 125 ms by using a DWI-EPI sequence. For diffusion encoding, we used the isotropic generalized diffusion tensor MRI (IGDTI) acquisition protocol to achieve an efficient orientationally averaged DW signal[3] with the following parameters: 16 linearly sampled *b*-values ranging from 2,540 to 14,700 s/mm$^2$ in 3 directions, 14 linearly sampled *b*-values ranging from 4,140 to 14,700 s/mm$^2$ in 4 directions, and 9 linearly sampled *b*-values ranging from 8,260 to 14,700 s/mm$^2$ in 6 directions, using the efficient gradient sampling schemes in Table 2 in Avram *et al.*[3] This type of diffusion encoding increases the contrast given by local anisotropy and is not intended to measure the isotropic diffusion in the system. Additional diffusion parameters were gradient duration of $\delta$=4 ms and diffusion time of $\Delta$=15 ms.

The 2D distribution of MD-$T_2$ was estimated (in conjunction with the *a priori* obtained 1D distributions as constraints) from a 2D D-$T_2$-weighted data set with 16 sampled combinations of echo times and *b*-values within the 1D acquisition range.

The data were averaged twice to maintain high signal-to-noise ratio (SNR), which was always maintained above 100 (defined as the ratio between the average unattenuated signal intensity within a tissue region of interest, and the standard deviation of the signal intensity within the background). The sample temperature was set at 16.8°C.

#### Harmonization of distributions between subjects

Since amyloid plaques, fibrillary tangles and microglial activation were less likely to affect white matter than gray matter, we chose white matter to be used as a reference region in each brain sample. White matter regions were manually delineated on each MRI slice with corresponding histological data (Supplementary Fig. 1A). To further restrict the reference MRI voxels to those least likely to be influenced by pathology, we defined a voxel-wise pathology score. Because pTau and microglial histology were available for the same slices used for MD-MRI, pTau and microglial histology values were Z-scored across the white matter region within each subject. The pathology score for each voxel was then calculated as the sum of the Z-scored pTau and microglial values (Supplementary Fig. 1B). For each subject, the 250 voxels with the lowest pathology scores were selected, and the mean T2–D distribution across these voxels was computed. This mean distribution served as the subject-specific reference distribution. The reference distribution from the control subject, computed using all white matter voxels, was used as a common reference to which all subject-specific reference distributions were aligned (Supplementary Fig. 1C).

To align each subject's reference distribution to the common reference spectrum, we first identified the region surrounding the highest peak (mode) of the reference distribution. This region was defined by applying a Gaussian filter (full width at half maximum = 3 grid points) to the spectrum, identifying grid points with intensities exceeding 50% of the peak value, and retaining only those grid points connected to the peak region. The mean grid coordinate of this region was then computed and defined as the reference point of the spectrum (Supplementary Fig. 1D). The difference in grid coordinate between a subject's reference point and the common reference point defined the shift required to align the subject-specific reference distribution to the common reference distribution (Supplementary Fig. 1E, F). This shift was applied to the distribution at every voxel—including non–white matter voxels—using modified Akima cubic Hermite interpolation (MATLAB function interp2).

#### Region-of-interest analysis of 3D virtual histology images

To generate the white matter ROIs, we first computed a mean diffusivity image from the MD-MRI distributions for each subject. We computed a histogram of the mean diffusivity values for each subject. After the application of Gaussian smoothing to the histograms, they were bimodal. We set a threshold at halfway between the two peaks and labeled voxels with mean diffusivity values lower than the threshold as white matter. Then, we performed the following operations to refine the white matter ROIs: fill holes (MATLAB command “imfill”), remove thin regions (MATLAB command “imopen” with a structural element “disk” 2 voxels wide as input), remove connected components that contain less than 300 voxels (connected components found with MATLAB command “bwconncomp”), erode mask by one layer of pixels (MATLAB command “imerode”), and remove the first 10 and last 10 slices because the slices at the edges of the volume had lower signal-to-noise ratios. After the white matter masks were created using those automated steps, they were manually refined to reduce over- and under-segmentation.

### Supplementary Figures

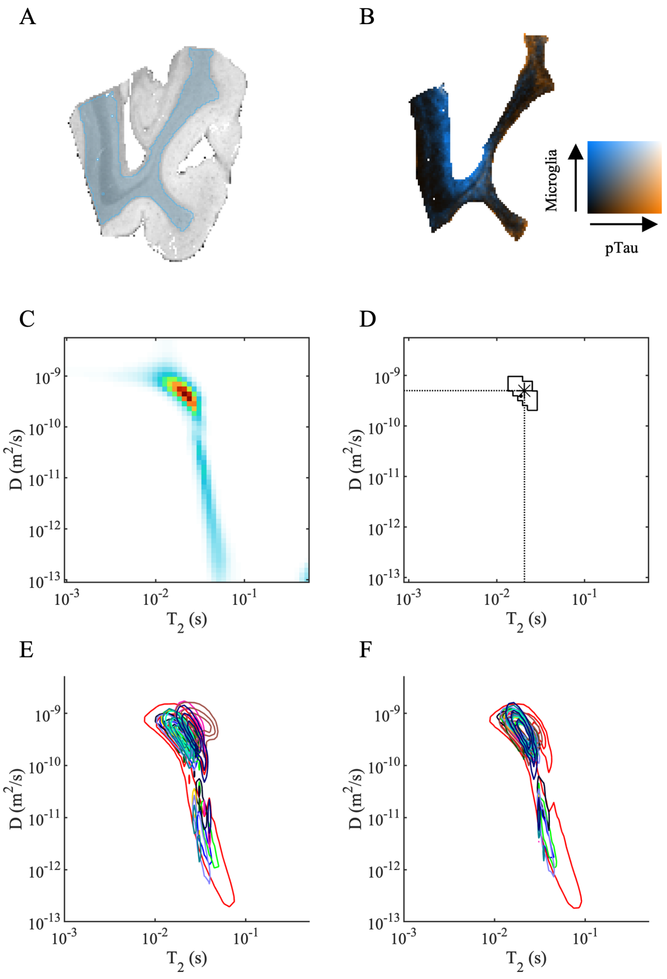


**Supplementary Figure 1. Harmonization of estimated $T_2$-D distributions between subjects. (A)** For each subject, a white matter (WM) region of interest (ROI) was manually outlined (blue). **(B)** For each WM voxel, a pathology score was determined as the sum of the Z-scored histological intensities of pTau and microglia. Black color indicates no pathology, orange color a high pTau intensity, blue color a high microglia intensity, and white a high combined pathology. **(C)** The mean $T_2$-D distribution over the WM ROI was determined for each of the subjects. **(D)** From the mean distribution, the extent of the peak region and its center coordinate was determined for each of the subjects. For each subject, the $T_2$-D distribution at each voxel was then translated to align with the control. The mean $T_2$-D distribution in the WM ROI shown for each subject in different color **(E)** before harmonization and **(F)** after harmonization.

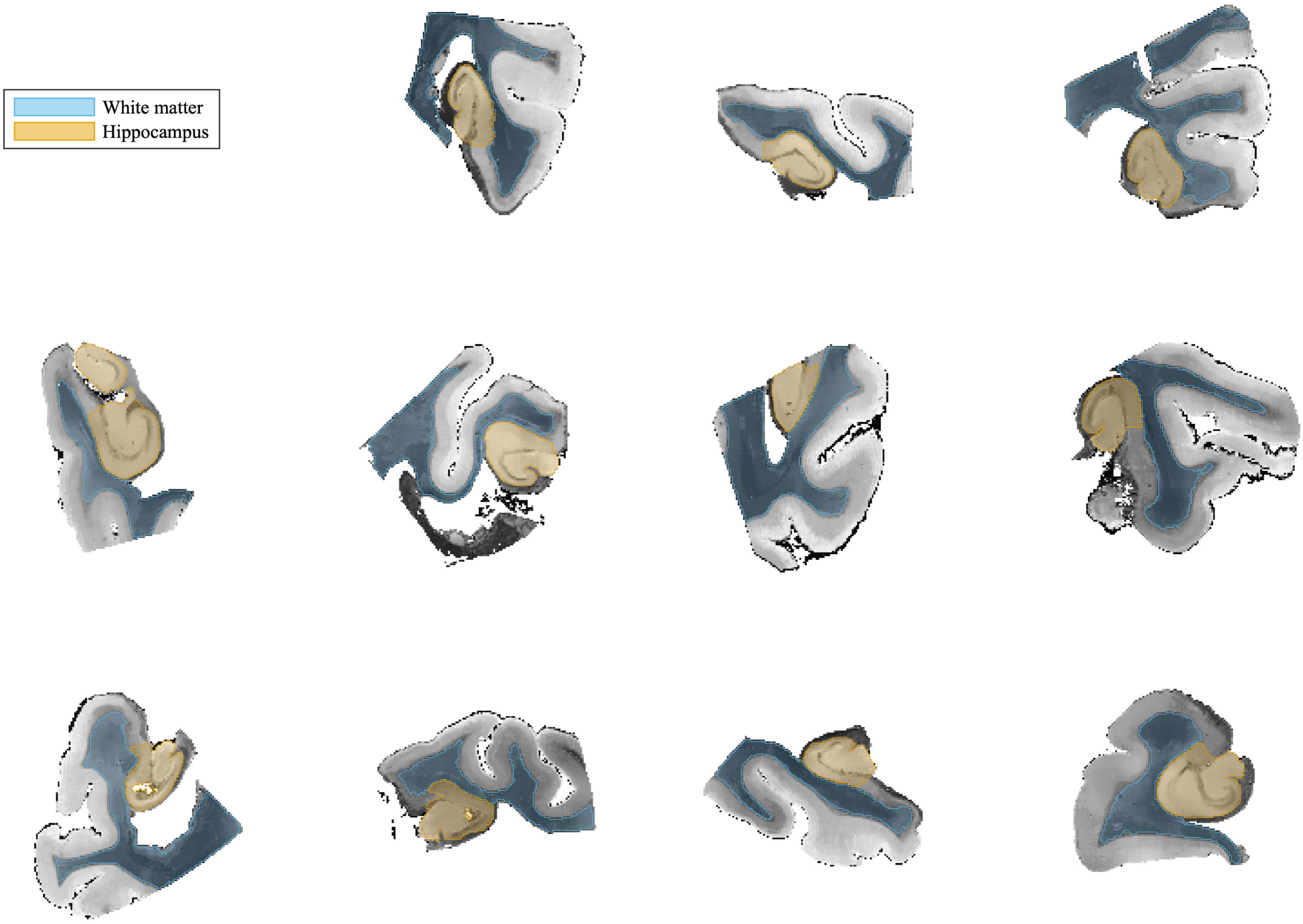


**Supplementary Figure 2. White matter and hippocampus regions of interest (ROI) used in the analysis.** White matter (blue) and the hippocampus (orange) ROIs were outlined for the analysis. Only one slice is shown for each subject. The hippocampus was further delineated into subfields which are shown in Supplementary Fig. 3.

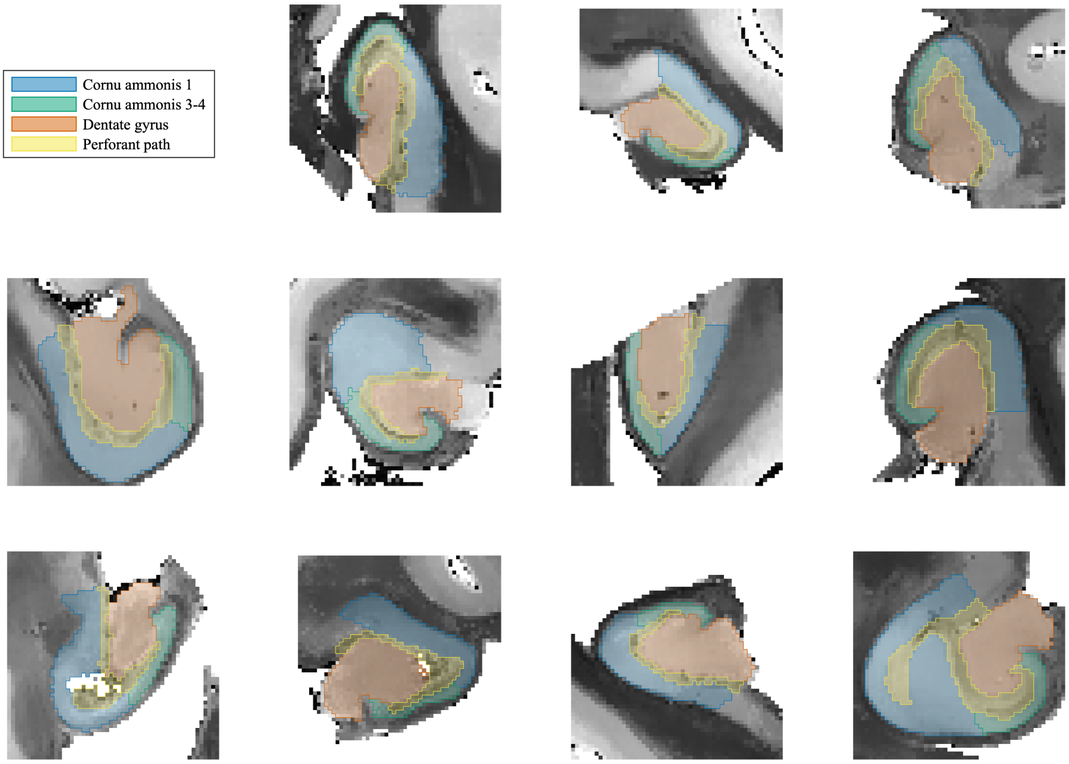


**Supplementary Figure 3. Hippocampus subfield regions of interest (ROI).** The hippocampus was delineated into the subfields cornu ammonis 1 (blue), cornu ammonis 3-4 (green), dentate gyrus (vermilion) and perforant path (yellow).

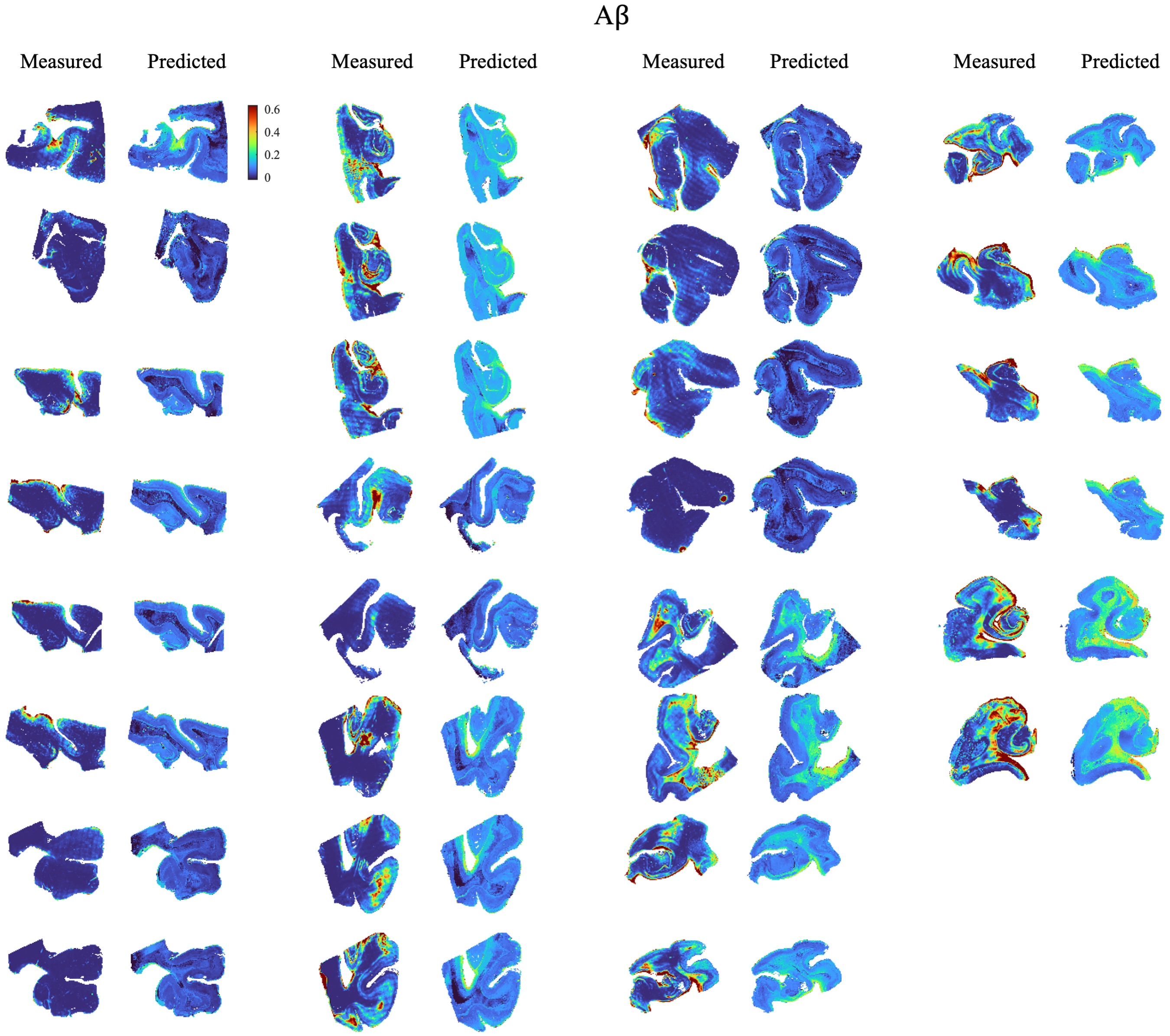


**Supplementary Figure 4. Measured and predicted A$\beta$ histological images.** Between one and four histological slices were available from each of the 12 subjects, resulting in a total of 30 slices.

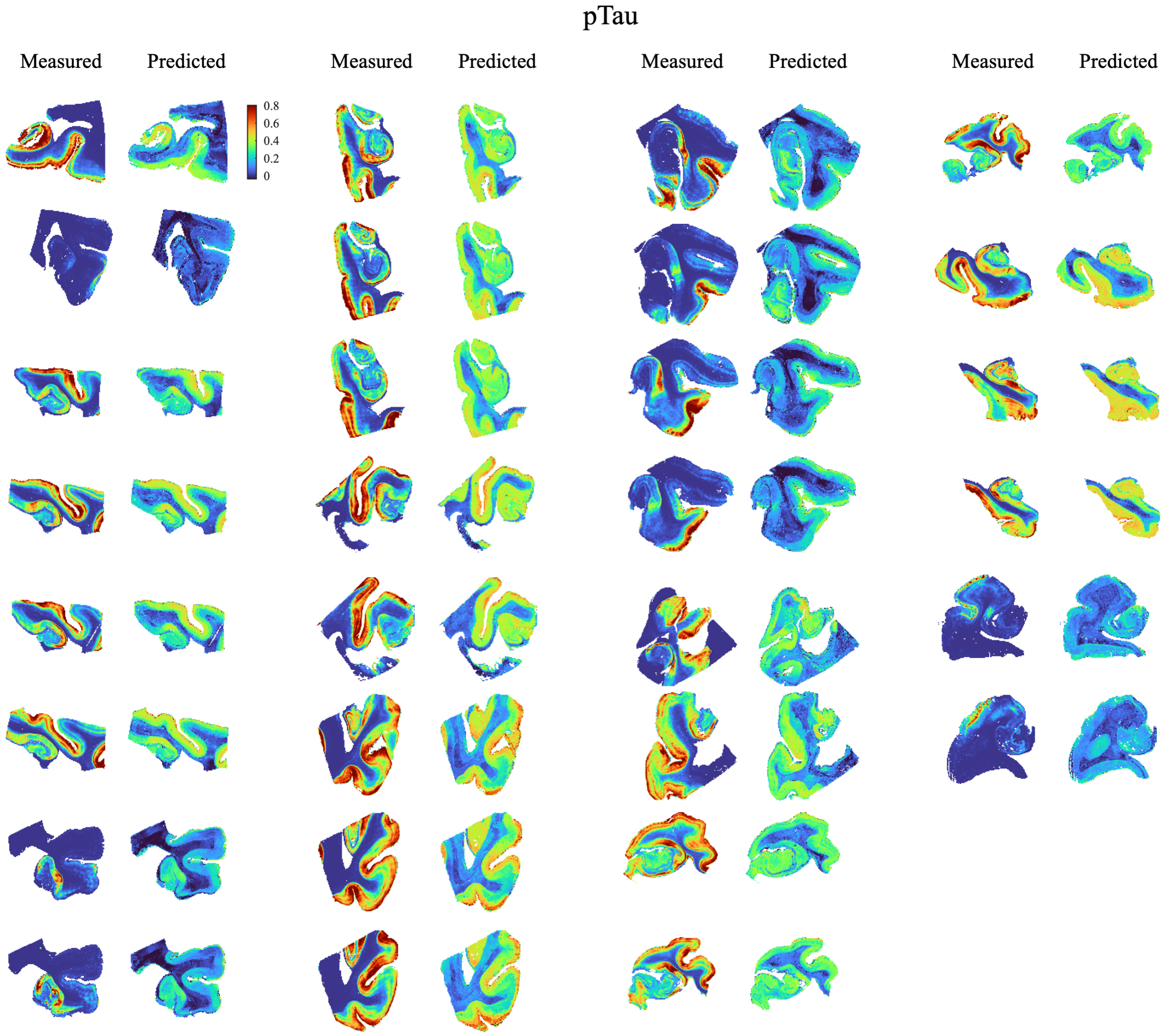


**Supplementary Figure 5. Measured and predicted pTau histological images.** Between one and four histological slices were available from each of the 12 subjects, resulting in a total of 30 slices.

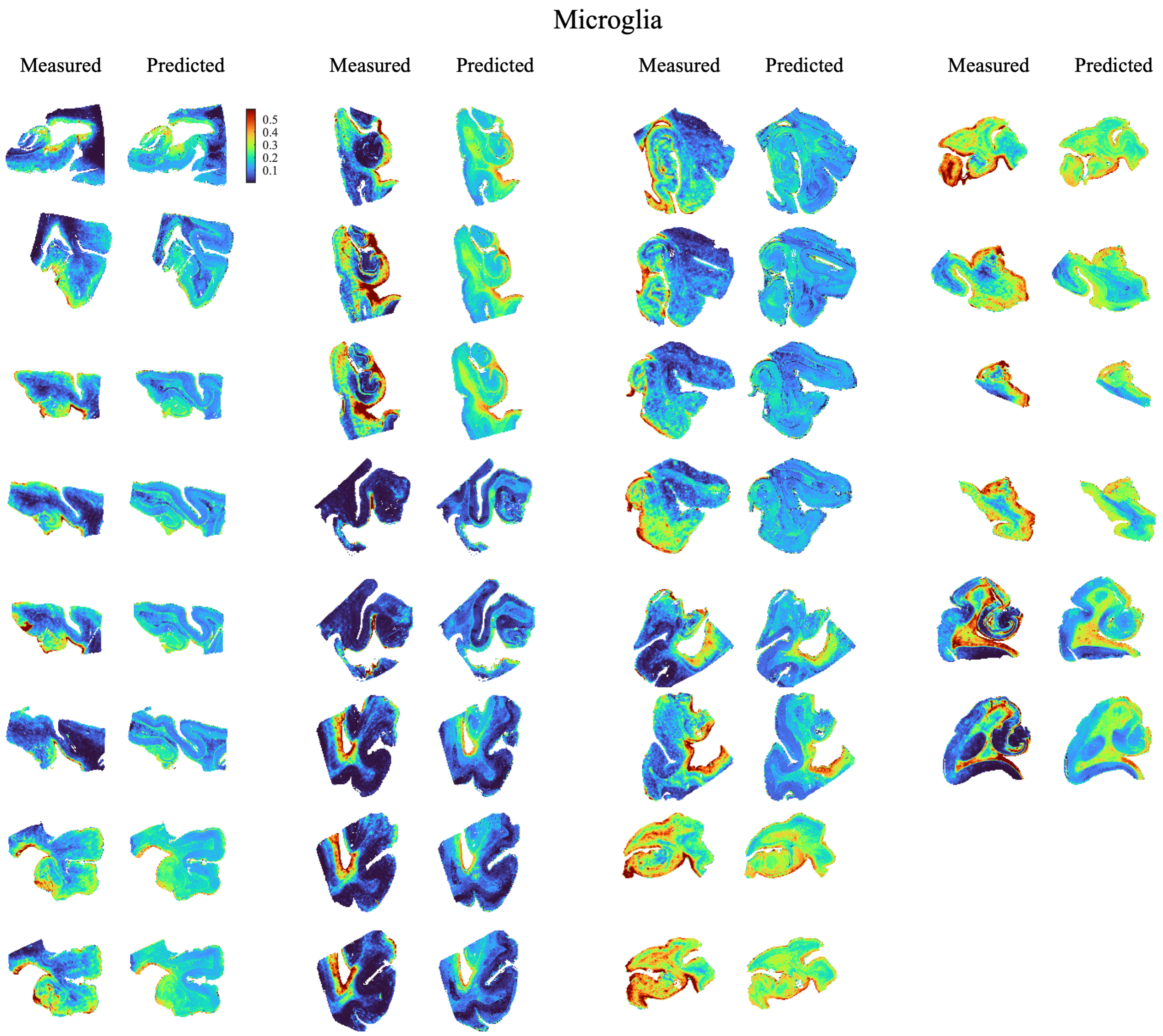


**Supplementary Figure 6. Measured and predicted microglia histological images.** Between one and four histological slices were available from each of the 12 subjects, resulting in a total of 30 slices.

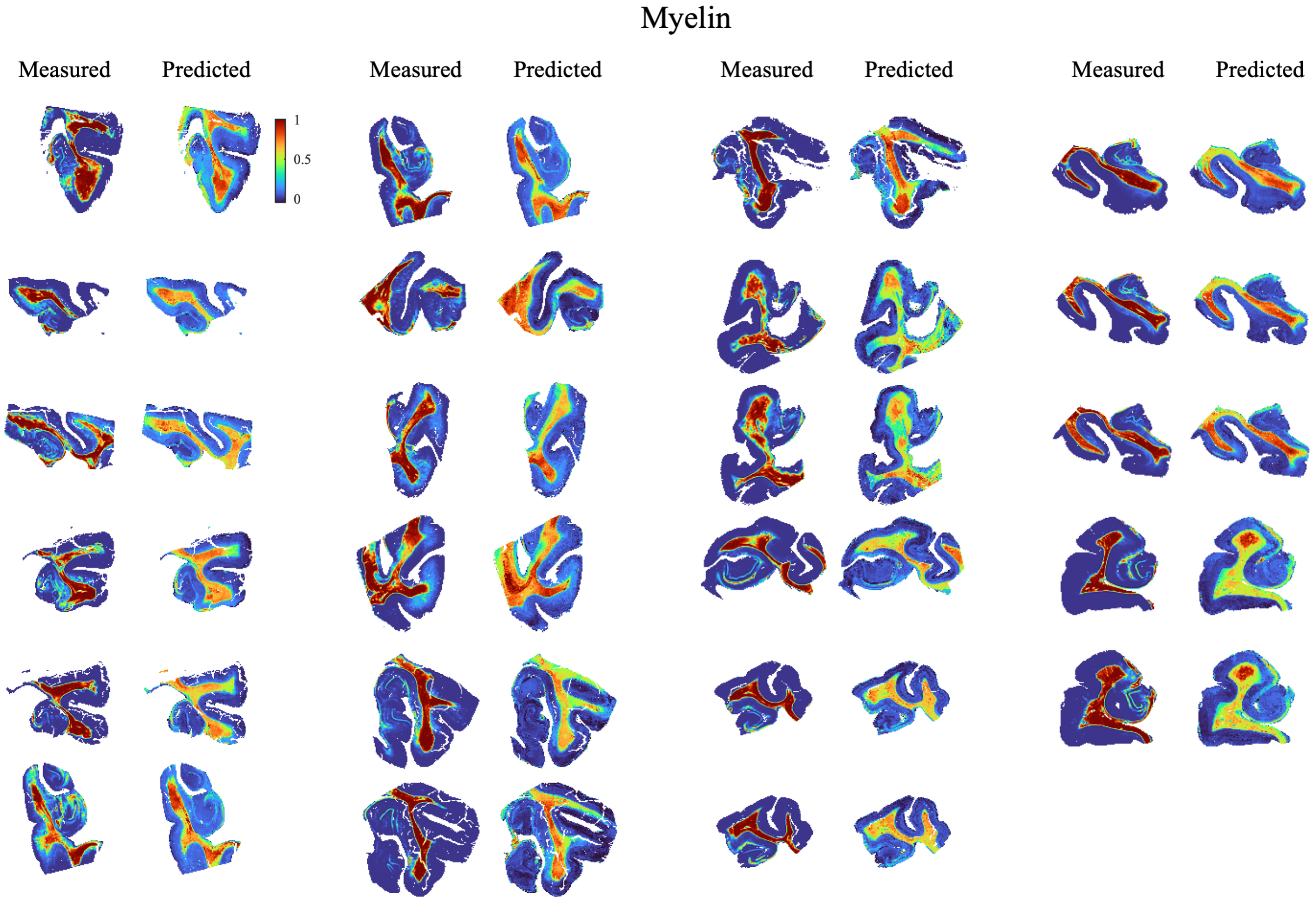


**Supplementary Figure 7. Measured and predicted myelin histological images.** Between one and three histological slices were available from each of the 11 subjects, resulting in a total of 23 slices.
